# Empirical evidence of the neuroactive potential of the gut on neurochemistry in the gut-brain axis in humans

**DOI:** 10.1101/2025.05.07.652653

**Authors:** Nicola Johnstone, Kathrin Cohen Kadosh

## Abstract

The gut microbiome produces a wide range of neuroactive metabolites capable of influencing brain neurotransmitter systems. While preclinical studies suggest these microbial pathways modulate cognitive and emotional processes, evidence in humans remains limited. This study investigates the relationship between gut microbiome-derived neuroactive pathways and in vivo brain neurotransmitter concentrations in healthy young females. Using proton magnetic resonance spectroscopy (¹H-MRS), we quantified GABA and glutamate levels in the dorsolateral prefrontal cortex (dlPFC), anterior cingulate cortex (ACC), and inferior occipital gyrus (IOG). Parallel gut microbiome profiling identified functional pathways associated with the synthesis and degradation of GABA, glutamate, short-chain fatty acids, p-cresol, and inositol. We observed significant, region-specific associations between microbial neuroactive pathways and both neurotransmitter concentrations and excitatory/inhibitory (E/I) balance, a key regulator of neuroplasticity and mental health. Notably, microbial GABA synthesis pathways were inversely related to brain glutamate levels in the dlPFC, while glutamate synthesis pathways correlated with GABA levels in the ACC. Additional pathways, including p-cresol and SCFA metabolism, showed region-dependent relationships with brain neurochemistry and E/I ratios. These neurochemical findings were complemented by associations between the same microbial pathways and psychological outcomes, including anxiety, depressive symptoms, and sleep quality. These findings provide new evidence of gut-derived metabolic influence on human brain neurotransmitter dynamics and mental wellbeing, highlighting the gut-brain axis as a promising target for microbiome-based interventions supporting cognitive resilience and emotional health.

## Introduction

A wealth of preclinical evidence has demonstrated strong links between the gut microbiome and brain function showing that gut microbial composition influences cognition, behaviour, and mental health [1–3]. Several mechanisms have been proposed to explain these effects, notably the microbiota-mediated production of neuroactive compounds, including gamma-aminobutyric acid (GABA), glutamate, serotonin, dopamine, tryptophan, and short-chain fatty acids (SCFAs, e.g., propionate, butyrate, acetate) [4–6]. These metabolites play critical roles in mood regulation, cognitive function, and neurodevelopmental disorders [3, 7–11], in part through modulation of cortical excitatory/inhibitory (E/I) balance—a key regulator of neuroplasticity, mental health, and cognition [11–13]. Additionally, gut-derived metabolites such as p-cresol have been implicated in modulating neuroplasticity and oxidative stress [14, 15], potentially influencing neurotransmitter dynamics and overall brain function [16].

Despite substantial preclinical evidence defining these gut-derived neuroactive mechanisms [17, 18], translating these findings into human studies remains challenging. Bridging this gap requires rigorous investigation into how microbial metabolites exert functionally relevant effects on human cognition, behaviour, and neurochemistry. Recent human studies have begun employing neuroimaging and metabolomics to investigate these relationships [4, 19], however, demonstrating causality and clinical significance remains challenging. Progress in this area demands well-designed human studies integrating multi-modal methodologies to elucidate gut-brain interactions and their therapeutic potential. Such approaches are particularly needed to assess whether gut microbiome activity influences both brain neurochemistry and psychological outcomes, including anxiety, mood, and sleep quality.

The current study aims to characterise the relationship between the neuroactive capacity of the gut microbiome and brain neurochemistry, specifically focusing on GABA and glutamate—key inhibitory and excitatory neurotransmitters. We hypothesise that gut-derived neuroactive pathways significantly correlate with region-specific GABA and glutamate concentrations, influencing cortical E/I balance. Understanding these interactions represents a critical step toward developing microbiome-based interventions for cognition, mood, and stress-related disorders. To achieve this, we aim to identify physiological signatures of microbial influence on brain function that may be adaptive or maladaptive, even within healthy individuals.

The gut microbiome plays a pivotal role in neurodevelopment, neural homeostasis, and behaviour [20–22]. Communication between the gut and brain is bidirectional, with neural signals regulating microbial activity and gut-derived metabolites modulating brain function via multiple pathways [23]. One key pathway for gut-brain signalling involves vagal transmission, where gut-derived metabolites engage the vagus nerve to relay information to the brain [24]. These metabolites, once absorbed systemically, can influence the integrity of both intestinal and blood-brain barriers (BBB) [25]. Although preclinical evidence suggests some microbial metabolites cross these barriers to affect central neurotransmitter systems, their direct human effects remain poorly characterised (ref). Additionally, certain microbial communities operate independently from central nervous regulation, producing bioactive metabolites that directly influence host physiology [26].

To systematically explore gut-brain interactions, recent studies have structured the microbially derived neuroactive potential into gut-brain modules (GBMs), using a library encompassing 56 metabolic pathways associated with neuroactive compound synthesis and degradation [19, 27]. GABA and glutamate metabolism pathways within these GBMs are of particular interest due to their involvement in neuropsychiatric disorders [5, 11, 28]. While gut-derived GABA is predicted to cross intestinal and blood-brain barriers [29], uncertainty remains regarding glutamate permeability due to predictive model limitations [30]. This study aims to address this knowledge gap by directly linking gut-derived neuroactive signatures with in vivo brain neurochemistry, focusing on indirect gut-brain signalling pathways.

Furthermore, mounting evidence suggests the gut microbiome significantly influences circulating metabolites, thereby impacting neurotransmitter balance and brain function. For example, gut microbiota composition has been associated with altered brain insulin sensitivity and corresponding shifts in metabolite profiles in mice [31]. Computational models have predicted that specific gut-derived metabolites can traverse biological barriers and influence neurotransmission [4, 19, 32]. Supporting this gut-metabolite-brain axis, alterations in gut microbial composition correlate with neurotransmitter disruptions in various neurodevelopmental and psychiatric disorders. Although our current study focuses on healthy human subjects, insights from clinical and preclinical studies of pathological conditions illustrate key mechanisms that may affect brain function broadly. For instance, preclinical models of autism spectrum disorder demonstrate that gut dysbiosis affects brain E/I balance by disrupting intestinal amino acid transport—a process reversible via targeted microbial supplementation [33].

However, determining causality remains challenging, as psychiatric conditions and associated medications (e.g., antipsychotics) can themselves induce gut dysbiosis, complicating interpretations of microbiota changes as cause or consequence of disease [34, 35]. Similarly, human studies of bipolar depression show correlations between functional connectivity patterns and gut-derived serum metabolites [8]. Employing a multi-omics approach, Wu et al. identified specific microbial taxa associated with circulating metabolites linked to altered brain network activity. Such findings further reinforce the hypothesis that gut microbiota significantly contribute to neuropsychiatric pathophysiology [36, 37]. Beyond psychiatric conditions, evidence from non-psychiatric health challenges such as metabolic syndrome and chronic pain conditions also illustrates how gut-derived metabolites influence brain function and symptomatology, underscoring the broader relevance of the gut-metabolite-brain axis [38].

Proton magnetic resonance spectroscopy (¹H-MRS) enables the quantification of human brain GABA and glutamate concentrations. Although ¹H-MRS does not directly measure synaptic neurotransmission, it provides valuable estimates of tonic neurochemical levels across intracellular and extracellular compartments, thereby reflecting underlying neural activity and cortical responsiveness [39]. By integrating these ¹H-MRS-derived brain measures with gut microbiome-associated metabolic signatures, the current study aims to elucidate the relationship between gut-derived metabolites and brain neurotransmitter dynamics. This integrative approach offers crucial insights toward identifying microbial pathways that may contribute to E/I balance and psychological wellbeing and represents an important step toward developing targeted microbiome-based therapeutic strategies.

## Methods

Ethical approval was obtained from the University ethics committee. All participants provided written informed consent, with additional consent from primary caregivers for participants under 18. Participants under 18 also provided assent before MRI procedures. Participants were compensated £75 for their participation.

### Participant characteristics and sample overview

A total of 83 healthy females aged 17–25 years (Mean = 20.32, SD = 2.62) participated in this double-blind, placebo-controlled trial. For the analyses presented here, baseline (pre-intervention) data across the entire sample were used (see for a full report on the trial outcomes [40]). Neurochemical data were successfully acquired via ¹H-MRS for 76 participants in the dorsolateral prefrontal cortex (DLPFC), 74 in the anterior cingulate cortex (ACC), and 72 in the inferior occipital gyrus (IOG), with slight variability due to exclusion criteria for spectral quality. Shotgun metagenomic sequencing of stool samples was successfully completed for 61 participants, with all samples collected prior to neuroimaging sessions. Eligibility criteria excluded individuals with psychiatric diagnoses, gastrointestinal conditions, restrictive diets, or MRI contraindications, thereby minimizing known confounders for gut-brain interaction studies.

### Procedure and materials

Participants underwent MRI scans lasting approximately one hour, comprising anatomical imaging and three ¹H-MRS scans. Volumes of interest (VOIs) were defined in three regions: left dorsolateral prefrontal cortex (dlPFC), anterior cingulate cortex (ACC), and left inferior occipital gyrus (IOG) [40]. These VOIs were selected based on their respective roles in emotional behaviour (dlPFC) [41, 42], cognitive control and regulation (ACC) [43, 44], and as a neutral reference region associated with visual processing (IOG)[45]. Participants collected stool samples at home using provided kits for subsequent microbiome analysis.

### MRI scanning

Anatomical acquisition involved using a Siemens 3T Magneton TIM Trio scanner with a 32-channel head coil. T1-weighted MPRAGE images (TR = 1900 ms, TE = 3030 ms, dwell time = 1500 ms, 1 mm slices, FOV = 256 x 256) were obtained in 5 minutes. T2*-weighted TSE images (TR = 3500 ms, TE = 93 ms, dwell time = 7100 ms, GRAPPA = 2, FOV = 179 x 256, 4 mm slices) were captured in the coronal and axial planes. Structural images were reconstructed for voxel placement in 1H-MRS acquisition. ¹H-MRS data were acquired from three volumes of interest (VOIs) in the left dorsolateral prefrontal cortex (DLPFC), anterior cingulate cortex (ACC), and left inferior occipital gyrus (IOG). A 2 cm³ voxel was manually centred within each VOI using sagittal T1-weighted images. To minimize inter-subject variability in voxel placement, anatomical landmarks were consistently used for voxel positioning across participants, and placement was guided by standardized neuroanatomical criteria and cross-referenced across sagittal, coronal, and axial planes. All placements were performed by the same trained operator and visually inspected by a second rater to ensure consistency.

Spectra acquisition used the Spin Echo full Intensity-Acquired Localized spectroscopy sequence, SPECIAL[46]. Water-suppressed spectra were collected with TR = 3200 ms, TE = 8.5 ms, and flip angle = 90° over 192 averages. Water-unsuppressed spectra were acquired beforehand over 16 averages in the same region. Saturation bands were aligned along the six outer planes of each voxel to minimise unwanted signals. **Figure 1** illustrates the voxel placement in each region of interest.

**Figure 1.**
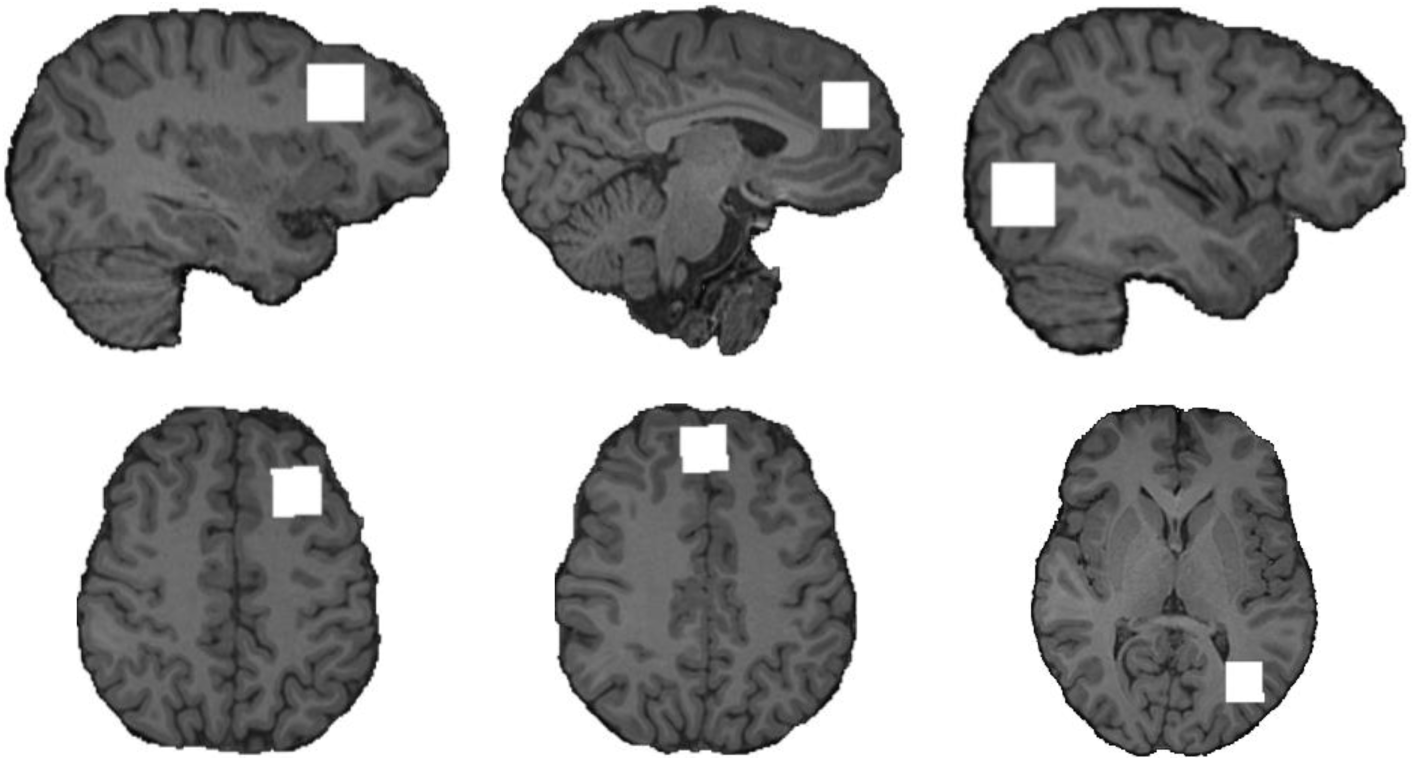
A representation of ¹H-MRS voxel locations over each region, shown as white squares, all 2 cm^3^. The most lateral sagittal edge and superior axial edge are shown (from left to right) for the dorsolateral prefrontal cortex VOI, anterior cingulate gyrus VOI, and inferior occipital gyrus VOI. Voxel locations were based on an anatomical reference image by the same trained operator and visually inspected by a second rater to ensure consistency.

¹H-MRS data pre-processing involved using FID-A toolbox functions in MATLAB (The Mathworks, Natick, MA) to remove motion-corrupted averages and correct frequency drift. Voxel tissue was apportioned for cerebrospinal fluid (CSF) correction [47] using the segment function of the Gannet toolbox [48] which uses the Statistical Parametric Mapping toolbox (SPM12, MATLAB, The Mathworks, Natick, MA) for co-registration and segmentation[49]. Detailed formulas for tissue correction are available at <u>Gannet documentation</u>.

Cleaned averaged spectra were analysed in LCModel [50] to estimate relative concentrations of GABA and glutamate in each VOI, referenced to the water peak. Spectral quality was assessed through: (i) line width of the water peak (full width at half maximum), excluding widths >13 Hz; (ii) signal-to-noise ratio (SNR), defined as the NAA peak height (1.8–2.2 ppm) divided by noise (averaged signal from −2 to 0 ppm), with SNR <150 excluded; (iii) visual confirmation of NAA, Cr, Cho, and glutamate peaks at expected ppm values; and (iv) Cramer-Rao lower bounds (CRLBs), with thresholds of >29 for GABA and >15 for glutamate. These stringent criteria ensured consistency across timepoints and VOIs.

### Gut microbiome

#### Stool sampling

Sampling kits were provided by BaseClear, the Netherlands including hygienic tools, stool catcher, collection tubes containing DNA/RNA Shield (Zymo Research, CA, USA) and secure return packaging with instructions. Samples were collected at home and kept at ambient room temperature until returned to the laboratory (within 1 month) for storage at −L80 °C.

#### Next generation sequencing: shallow shotgun metagenomics

All samples were shipped to Novogene, Cambridge, UK, for shallow shotgun metagenomic analysis[51]. This process involved several stages, including DNA extraction, quality control (QC), metagenomic library preparation, sequencing, and subsequent taxonomic and functional annotation. Samples that met QC thresholds, which included absence of contamination and sufficient volume, proceeded to library construction.

The library construction process involved randomly fragmenting genomic DNA into approximately 350 bp segments using a Covaris ultrasonic disruptor. These fragments were then end-repaired, A-tailed, and ligated with sequencing adapters. After this, the library was size-selected, PCR-amplified, purified, and quantified using Qubit and qPCR methods.

Libraries that passed QC, including those with effective library concentrations exceeding 3 nM, were pooled according to target data output requirements, and subjected to shallow shotgun sequencing. Sequencing was performed on the NovaSeq X Plus Series platform, using paired-end 150 bp reads (PE150), and generated 6 GB of raw data per sample.

After sequencing, quality control was conducted using Fastp [52, 53] to remove paired reads that contained adapter contamination, had more than 10% uncertain nucleotides, or contained more than 50% low-quality nucleotides (base quality <5). To further ensure data quality, clean reads were aligned to the host genome using Bowtie2 [54] with the parameters--end-to-end,--sensitive, -I 200, -X 400, following previously established protocols [55–57].

The filtered clean data was assembled into contigs using MEGAHIT with meta-large presets [58]. During the assembly process, scaftigs, which are scaffolds without ‘N’ regions, were produced by breaking scaffolds at the ‘N’ junctions[59]. Gene prediction was carried out using MetaGeneMark on scaffolds greater than or equal to 500 bp in length [60], and sequences shorter than 100 nucleotides were filtered out[59]. To remove redundancy, the predicted genes were dereplicated using CD-HIT, creating a non-redundant gene catalogue [61]. For taxonomic annotation, the predicted genes were aligned to the MicroNR database (https://www.ncbi.nlm.nih.gov), which contains sequences from bacteria, fungi, archaea, and viruses. The alignment was performed using DIAMOND [62], employing the blastp algorithm with parameters set to -e 1e-5 [55]. The taxonomy of the sequences was assigned using the lowest common ancestor (LCA) algorithm, implemented in MEGAN [63]. Species abundance was calculated by summing the gene abundances corresponding to each species [57, 64, 65].

Function annotation was inferred from KEGG ORTHOLOGY (KO) database, which represents molecular functions as functional orthologs. Unigenes were compared using DIAMOND software (blastp, eval <= 1e-5). For each sequence, the result with the highest score (one HSP > 60 bits) was selected for analysis. The relative abundance was calculated as the sum of the relative abundance of the genes annotated. For a given function, the number of genes in a sample is equal to the number of genes with non-zero abundance among those annotated for that function.

#### Bioinformatic analysis of sequenced stool samples

Gut-brain modules (GBMs) were identified using Omixer-RPM in R [66]. To ensure sufficient representation of a module at the group level, a threshold of (minimum 30% coverage was applied, with 10% prevalence [67]. Functional relationships between the gut and brain were analysed using the R package Microbiome Multivariable Association with Linear Models[68], MaAsLin2 (version 1.15.1). We assessed the association of GABA and Glutamate in each brain region (ACC, dlPFC, IOG) with neuroactive pathways in one model and the E/I ratio in another.

To identify neuroactive pathways reliably linked to neurochemical levels, we calculated Storey’s q-values alongside MaAsLin2’s p- and q-values to control for the local false discovery rate (lfdr) following established best practices for multi-omics association analysis [68]. While MaAsLin2 employs the Benjamini-Hochberg (BH) method for FDR control, Storey’s approach estimates the proportion of true null hypotheses, providing a more adaptive and data-driven correction. Integrating both methods enhances statistical robustness, balancing conservative and adaptive FDR control for reliable findings.

We applied Leave-One-Out (LOO) cross-validation with MaAsLin2 to assess result consistency. In each iteration, one sample was excluded, the model rerun, and coefficients and p-values aggregated. 95% confidence intervals were calculated from aggregate data. Confidence intervals that do not cross zero are considered true effects and are reported.

## Results

From 61 sequenced stool samples, 23 GBMs met the inclusion criteria. (**Table 1**).

**Table 1.** Strength and direction of relationship between the neurochemicals Glutamate and GABA by region and neuroactive pathways of gut microbiota.

| Pathway Function | Neuroactive Pathway | Predictor | Region | coef | stderr | N | N.r | p | q | lfr | coi | conf_lc | conf_hi | Effect |
| --- | --- | --- | --- | --- | --- | --- | --- | --- | --- | --- | --- | --- | --- | --- |
| Amino Acid Metabolism | GABA degradation | GABA | ACC | 0.046 | 0.122 | 60 | 60 | 0.709 | 0.927 | 0.976 | 0 | -0.193 | 0.286 | NA |
| Amino Acid Metabolism | GABA degradation | GABA | DLPFC | 0.067 | 0.102 | 60 | 60 | 0.519 | 0.845 | 0.975 | 0 | -0.134 | 0.267 | NA |
| Amino Acid Metabolism | GABA degradation | GABA | IOG | 0.111 | 0.205 | 60 | 60 | 0.595 | 0.892 | 0.976 | 0 | -0.292 | 0.513 | NA |
| Amino Acid Metabolism | GABA degradation | Glutamate | ACC | -0.108 | 0.124 | 60 | 60 | 0.395 | 0.780 | 0.953 | 0 | -0.351 | 0.135 | NA |
| Amino Acid Metabolism | GABA degradation | Glutamate | DLPFC | -0.143 | 0.122 | 60 | 60 | 0.254 | 0.702 | 0.868 | 0 | -0.382 | 0.097 | NA |
| Amino Acid Metabolism | GABA degradation | Glutamate | IOG | -0.113 | 0.128 | 60 | 60 | 0.383 | 0.775 | 0.951 | 0 | -0.363 | 0.137 | NA |
| Amino Acid Metabolism | GABA synthesis I | GABA | ACC | -1.346 | 1.231 | 60 | 29 | 0.284 | 0.718 | 0.904 | 0 | -3.759 | 1.066 | NA |
| Amino Acid Metabolism | GABA synthesis I | GABA | DLPFC | -0.534 | 1.030 | 60 | 29 | 0.608 | 0.896 | 0.976 | 0 | -2.553 | 1.484 | NA |
| Amino Acid Metabolism | GABA synthesis I | GABA | IOG | -3.085 | 2.071 | 60 | 29 | 0.147 | 0.627 | 0.718 | 0 | -7.144 | 0.974 | NA |
| Amino Acid Metabolism | GABA synthesis I | Glutamate | ACC | 0.419 | 1.249 | 60 | 29 | 0.741 | 0.936 | 0.977 | 0 | -2.029 | 2.866 | NA |
| **Amino Acid Metabolism** | **GABA synthesis I** | **Glutamate** | **DLPFC** | **-3.482** | **1.232** | **60** | **29** | **0.008** | **0.475** | **0.380** | **60** | **-5.896** | **-1.068** | **TRUE** |
| Amino Acid Metabolism | GABA synthesis I | Glutamate | IOG | 0.253 | 1.285 | 60 | 29 | 0.839 | 0.956 | 0.977 | 0 | -2.266 | 2.772 | NA |
| Amino Acid Metabolism | GABA synthesis II | GABA | ACC | -2.233 | 1.278 | 60 | 37 | 0.091 | 0.614 | 0.615 | 3 | -4.738 | 0.272 | NA |
| Amino Acid Metabolism | GABA synthesis II | GABA | DLPFC | -0.271 | 1.070 | 60 | 37 | 0.803 | 0.950 | 0.977 | 0 | -2.367 | 1.826 | NA |
| **Amino Acid Metabolism** | **GABA synthesis II** | **GABA** | **IOG** | **-4.749** | **2.150** | **60** | **37** | **0.034** | **0.577** | **0.497** | **58** | **-8.963** | **-0.535** | **TRUE** |
| Amino Acid Metabolism | GABA synthesis II | Glutamate | ACC | 1.572 | 1.297 | 60 | 37 | 0.237 | 0.684 | 0.848 | 0 | -0.970 | 4.114 | NA |
| Amino Acid Metabolism | GABA synthesis II | Glutamate | DLPFC | -2.204 | 1.279 | 60 | 37 | 0.094 | 0.612 | 0.623 | 1 | -4.711 | 0.304 | NA |
| Amino Acid Metabolism | GABA synthesis II | Glutamate | IOG | 1.185 | 1.335 | 60 | 37 | 0.384 | 0.773 | 0.954 | 0 | -1.431 | 3.800 | NA |
| Amino Acid Metabolism | Glutamate degradation II | GABA | ACC | -0.626 | 0.573 | 60 | 58 | 0.283 | 0.720 | 0.914 | 0 | -1.750 | 0.497 | NA |
| Amino Acid Metabolism | Glutamate degradation II | GABA | DLPFC | -0.087 | 0.480 | 60 | 58 | 0.847 | 0.963 | 0.977 | 0 | -1.027 | 0.853 | NA |
| Amino Acid Metabolism | Glutamate degradation II | GABA | IOG | 0.356 | 0.964 | 60 | 58 | 0.708 | 0.928 | 0.974 | 0 | -1.534 | 2.245 | NA |
| Amino Acid Metabolism | Glutamate degradation II | Glutamate | ACC | 0.186 | 0.582 | 60 | 58 | 0.747 | 0.937 | 0.977 | 0 | -0.954 | 1.326 | NA |
| Amino Acid Metabolism | Glutamate degradation II | Glutamate | DLPFC | 0.494 | 0.574 | 60 | 58 | 0.397 | 0.779 | 0.959 | 0 | -0.630 | 1.619 | NA |
| Amino Acid Metabolism | Glutamate degradation II | Glutamate | IOG | 0.595 | 0.598 | 60 | 58 | 0.331 | 0.742 | 0.939 | 0 | -0.578 | 1.768 | NA |
| **Amino Acid Metabolism** | **Glutamate synthesis I** | **GABA** | **ACC** | **0.042** | **0.017** | **60** | **60** | **0.021** | **0.562** | **0.452** | **60** | **0.008** | **0.075** | **TRUE** |
| Amino Acid Metabolism | Glutamate synthesis I | GABA | DLPFC | 0.015 | 0.014 | 60 | 60 | 0.314 | 0.728 | 0.928 | 1 | -0.013 | 0.043 | NA |
| Amino Acid Metabolism | Glutamate synthesis I | GABA | IOG | -0.015 | 0.029 | 60 | 60 | 0.620 | 0.899 | 0.976 | 0 | -0.071 | 0.042 | NA |
| **Amino Acid Metabolism** | **Glutamate synthesis I** | **Glutamate** | **ACC** | **-0.036** | **0.017** | **60** | **60** | **0.050** | **0.599** | **0.533** | **41** | **-0.070** | **-0.001** | **TRUE** |
| Amino Acid Metabolism | Glutamate synthesis I | Glutamate | DLPFC | 0.023 | 0.017 | 60 | 60 | 0.193 | 0.636 | 0.772 | 0 | -0.011 | 0.057 | NA |
| Amino Acid Metabolism | Glutamate synthesis I | Glutamate | IOG | -0.013 | 0.018 | 60 | 60 | 0.493 | 0.833 | 0.966 | 0 | -0.048 | 0.023 | NA |
| Amino Acid Metabolism | Glutamate synthesis II | GABA | ACC | 0.019 | 0.022 | 60 | 60 | 0.409 | 0.786 | 0.958 | 0 | -0.025 | 0.062 | NA |
| Amino Acid Metabolism | Glutamate synthesis II | GABA | DLPFC | 0.030 | 0.018 | 60 | 60 | 0.112 | 0.618 | 0.647 | 0 | -0.006 | 0.067 | NA |
| Amino Acid Metabolism | Glutamate synthesis II | GABA | IOG | 0.003 | 0.037 | 60 | 60 | 0.909 | 0.976 | 0.977 | 0 | -0.070 | 0.075 | NA |
| Amino Acid Metabolism | Glutamate synthesis II | Glutamate | ACC | 0.009 | 0.022 | 60 | 60 | 0.700 | 0.926 | 0.976 | 0 | -0.035 | 0.053 | NA |
| Amino Acid Metabolism | Glutamate synthesis II | Glutamate | DLPFC | -0.016 | 0.022 | 60 | 60 | 0.470 | 0.815 | 0.966 | 0 | -0.059 | 0.027 | NA |
| Amino Acid Metabolism | Glutamate synthesis II | Glutamate | IOG | -0.041 | 0.023 | 60 | 60 | 0.082 | 0.611 | 0.604 | 2 | -0.087 | 0.004 | NA |
| Amino Acid Metabolism | Tryptophan degradation | GABA | ACC | 0.117 | 0.092 | 60 | 60 | 0.217 | 0.651 | 0.807 | 1 | -0.062 | 0.297 | NA |
| Amino Acid Metabolism | Tryptophan degradation | GABA | DLPFC | -0.102 | 0.077 | 60 | 60 | 0.193 | 0.634 | 0.773 | 0 | -0.253 | 0.048 | NA |
| Amino Acid Metabolism | Tryptophan degradation | GABA | IOG | 0.073 | 0.154 | 60 | 60 | 0.645 | 0.901 | 0.970 | 0 | -0.230 | 0.375 | NA |
| Amino Acid Metabolism | Tryptophan degradation | Glutamate | ACC | -0.127 | 0.093 | 60 | 60 | 0.186 | 0.633 | 0.772 | 0 | -0.309 | 0.056 | NA |
| Amino Acid Metabolism | Tryptophan degradation | Glutamate | DLPFC | 0.100 | 0.092 | 60 | 60 | 0.289 | 0.713 | 0.897 | 0 | -0.080 | 0.280 | NA |
| Amino Acid Metabolism | Tryptophan degradation | Glutamate | IOG | 0.031 | 0.096 | 60 | 60 | 0.746 | 0.939 | 0.976 | 0 | -0.157 | 0.218 | NA |
| Amino Acid Metabolism | Tryptophan synthesis | GABA | ACC | -0.211 | 0.204 | 60 | 60 | 0.311 | 0.731 | 0.927 | 0 | -0.611 | 0.189 | NA |
| Amino Acid Metabolism | Tryptophan synthesis | GABA | DLPFC | 0.052 | 0.171 | 60 | 60 | 0.754 | 0.939 | 0.977 | 0 | -0.283 | 0.386 | NA |
| Amino Acid Metabolism | Tryptophan synthesis | GABA | IOG | -0.021 | 0.343 | 60 | 60 | 0.930 | 0.981 | 0.977 | 0 | -0.694 | 0.652 | NA |
| Amino Acid Metabolism | Tryptophan synthesis | Glutamate | ACC | 0.006 | 0.207 | 60 | 60 | 0.932 | 0.979 | 0.977 | 0 | -0.400 | 0.412 | NA |
| Amino Acid Metabolism | Tryptophan synthesis | Glutamate | DLPFC | -0.015 | 0.204 | 60 | 60 | 0.931 | 0.981 | 0.977 | 0 | -0.415 | 0.386 | NA |
| Amino Acid Metabolism | Tryptophan synthesis | Glutamate | IOG | 0.231 | 0.213 | 60 | 60 | 0.288 | 0.725 | 0.908 | 0 | -0.186 | 0.649 | NA |
| Others | g Hydroxybutyric acid GHB degradation | GABA | ACC | 0.004 | 0.065 | 60 | 60 | 0.904 | 0.974 | 0.977 | 0 | -0.124 | 0.132 | NA |
| Others | g Hydroxybutyric acid GHB degradation | GABA | DLPFC | -0.003 | 0.055 | 60 | 60 | 0.925 | 0.976 | 0.977 | 0 | -0.110 | 0.104 | NA |
| Others | g Hydroxybutyric acid GHB degradation | GABA | IOG | 0.200 | 0.110 | 60 | 60 | 0.080 | 0.617 | 0.601 | 5 | -0.015 | 0.415 | NA |
| Others | g Hydroxybutyric acid GHB degradation | Glutamate | ACC | -0.036 | 0.066 | 60 | 60 | 0.588 | 0.889 | 0.974 | 0 | -0.166 | 0.093 | NA |
| Others | g Hydroxybutyric acid GHB degradation | Glutamate | DLPFC | -0.093 | 0.065 | 60 | 60 | 0.165 | 0.627 | 0.751 | 0 | -0.220 | 0.035 | NA |
| Others | g Hydroxybutyric acid GHB degradation | Glutamate | IOG | -0.007 | 0.068 | 60 | 60 | 0.879 | 0.970 | 0.977 | 0 | -0.141 | 0.126 | NA |
| Others | Inositol synthesis | GABA | ACC | 0.095 | 0.065 | 60 | 60 | 0.154 | 0.627 | 0.727 | 0 | -0.032 | 0.223 | NA |
| Others | Inositol synthesis | GABA | DLPFC | 0.018 | 0.054 | 60 | 60 | 0.744 | 0.939 | 0.977 | 0 | -0.089 | 0.124 | NA |
| **Others** | **Inositol synthesis** | **GABA** | **IOG** | **-0.317** | **0.109** | **60** | **60** | **0.006** | **0.476** | **0.366** | **60** | **-0.531** | **-0.102** | **TRUE** |
| Others | Inositol synthesis | Glutamate | ACC | 0.014 | 0.066 | 60 | 60 | 0.827 | 0.953 | 0.977 | 0 | -0.115 | 0.143 | NA |
| Others | Inositol synthesis | Glutamate | DLPFC | -0.005 | 0.065 | 60 | 60 | 0.927 | 0.981 | 0.977 | 0 | -0.132 | 0.123 | NA |
| Others | Inositol synthesis | Glutamate | IOG | 0.016 | 0.068 | 60 | 60 | 0.809 | 0.950 | 0.977 | 0 | -0.117 | 0.149 | NA |
| Others | Isovaleric acid synthesis I KADH pathway | GABA | ACC | 0.037 | 0.028 | 60 | 60 | 0.197 | 0.634 | 0.780 | 0 | -0.017 | 0.091 | NA |
| Others | Isovaleric acid synthesis I KADH pathway | GABA | DLPFC | -0.028 | 0.023 | 60 | 60 | 0.237 | 0.677 | 0.833 | 1 | -0.074 | 0.017 | NA |
| Others | Isovaleric acid synthesis I KADH pathway | GABA | IOG | 0.047 | 0.047 | 60 | 60 | 0.326 | 0.733 | 0.926 | 1 | -0.044 | 0.139 | NA |
| Others | Isovaleric acid synthesis I KADH pathway | Glutamate | ACC | -0.044 | 0.028 | 60 | 60 | 0.132 | 0.624 | 0.686 | 0 | -0.099 | 0.011 | NA |
| Others | Isovaleric acid synthesis I KADH pathway | Glutamate | DLPFC | 0.031 | 0.028 | 60 | 60 | 0.280 | 0.710 | 0.883 | 0 | -0.023 | 0.085 | NA |
| Others | Isovaleric acid synthesis I KADH pathway | Glutamate | IOG | 0.009 | 0.029 | 60 | 60 | 0.749 | 0.938 | 0.976 | 0 | -0.048 | 0.066 | NA |
| Others | Isovaleric acid synthesis II KADC pathway | GABA | ACC | 0.025 | 0.025 | 60 | 60 | 0.323 | 0.726 | 0.933 | 0 | -0.024 | 0.074 | NA |
| Others | Isovaleric acid synthesis II KADC pathway | GABA | DLPFC | -0.032 | 0.021 | 60 | 60 | 0.147 | 0.624 | 0.689 | 1 | -0.073 | 0.009 | NA |
| Others | Isovaleric acid synthesis II KADC pathway | GABA | IOG | 0.050 | 0.042 | 60 | 60 | 0.253 | 0.696 | 0.865 | 1 | -0.033 | 0.132 | NA |
| Others | Isovaleric acid synthesis II KADC pathway | Glutamate | ACC | -0.022 | 0.025 | 60 | 60 | 0.394 | 0.772 | 0.961 | 0 | -0.072 | 0.027 | NA |
| Others | Isovaleric acid synthesis II KADC pathway | Glutamate | DLPFC | 0.020 | 0.025 | 60 | 60 | 0.413 | 0.780 | 0.958 | 0 | -0.028 | 0.069 | NA |
| Others | Isovaleric acid synthesis II KADC pathway | Glutamate | IOG | -0.019 | 0.026 | 60 | 60 | 0.470 | 0.819 | 0.971 | 0 | -0.070 | 0.032 | NA |
| Others | Nitric oxide degradation II NO reductase | GABA | ACC | 0.629 | 0.764 | 60 | 55 | 0.419 | 0.790 | 0.956 | 0 | -0.869 | 2.127 | NA |
| Others | Nitric oxide degradation II NO reductase | GABA | DLPFC | -0.044 | 0.640 | 60 | 55 | 0.931 | 0.982 | 0.977 | 0 | -1.298 | 1.210 | NA |
| Others | Nitric oxide degradation II NO reductase | GABA | IOG | -0.341 | 1.286 | 60 | 55 | 0.789 | 0.946 | 0.976 | 0 | -2.862 | 2.180 | NA |
| Others | Nitric oxide degradation II NO reductase | Glutamate | ACC | 0.264 | 0.776 | 60 | 55 | 0.730 | 0.933 | 0.974 | 0 | -1.257 | 1.784 | NA |
| Others | Nitric oxide degradation II NO reductase | Glutamate | DLPFC | -0.016 | 0.765 | 60 | 55 | 0.959 | 0.984 | 0.977 | 0 | -1.515 | 1.484 | NA |
| Others | Nitric oxide degradation II NO reductase | Glutamate | IOG | 0.388 | 0.798 | 60 | 55 | 0.631 | 0.904 | 0.974 | 0 | -1.177 | 1.953 | NA |
| Others | p Cresol synthesis | GABA | ACC | 0.056 | 0.040 | 60 | 60 | 0.172 | 0.626 | 0.759 | 0 | -0.022 | 0.134 | NA |
| Others | p Cresol synthesis | GABA | DLPFC | 0.001 | 0.033 | 60 | 60 | 0.952 | 0.982 | 0.979 | 0 | -0.065 | 0.066 | NA |
| **Others** | **p Cresol synthesis** | **GABA** | **IOG** | **-0.174** | **0.067** | **60** | **60** | **0.015** | **0.522** | **0.423** | **59** | **-0.305** | **-0.043** | **TRUE** |
| Others | p Cresol synthesis | Glutamate | ACC | 0.010 | 0.040 | 60 | 60 | 0.803 | 0.951 | 0.978 | 0 | -0.069 | 0.089 | NA |
| Others | p Cresol synthesis | Glutamate | DLPFC | 0.001 | 0.040 | 60 | 60 | 0.949 | 0.984 | 0.979 | 0 | -0.077 | 0.079 | NA |
| Others | p Cresol synthesis | Glutamate | IOG | 0.002 | 0.042 | 60 | 60 | 0.919 | 0.975 | 0.978 | 0 | -0.079 | 0.084 | NA |
| Others | Quinolinic acid degradation | GABA | ACC | 0.034 | 0.018 | 60 | 60 | 0.070 | 0.607 | 0.574 | 6 | -0.001 | 0.070 | NA |
| Others | Quinolinic acid degradation | GABA | DLPFC | -0.011 | 0.015 | 60 | 60 | 0.466 | 0.818 | 0.969 | 0 | -0.041 | 0.019 | NA |
| Others | Quinolinic acid degradation | GABA | IOG | -0.024 | 0.031 | 60 | 60 | 0.440 | 0.799 | 0.965 | 0 | -0.084 | 0.036 | NA |
| Others | Quinolinic acid degradation | Glutamate | ACC | -0.027 | 0.019 | 60 | 60 | 0.159 | 0.628 | 0.734 | 0 | -0.063 | 0.009 | NA |
| Others | Quinolinic acid degradation | Glutamate | DLPFC | 0.032 | 0.018 | 60 | 60 | 0.093 | 0.619 | 0.603 | 1 | -0.003 | 0.068 | NA |
| Others | Quinolinic acid degradation | Glutamate | IOG | -0.014 | 0.019 | 60 | 60 | 0.489 | 0.832 | 0.967 | 0 | -0.051 | 0.024 | NA |
| SCFAs | Acetate degradation | GABA | ACC | 0.021 | 0.045 | 60 | 60 | 0.647 | 0.908 | 0.976 | 0 | -0.068 | 0.110 | NA |
| SCFAs | Acetate degradation | GABA | DLPFC | 0.034 | 0.038 | 60 | 60 | 0.383 | 0.775 | 0.950 | 0 | -0.041 | 0.108 | NA |
| SCFAs | Acetate degradation | GABA | IOG | -0.132 | 0.076 | 60 | 60 | 0.093 | 0.615 | 0.622 | 1 | -0.281 | 0.017 | NA |
| SCFAs | Acetate degradation | Glutamate | ACC | -0.010 | 0.046 | 60 | 60 | 0.815 | 0.949 | 0.977 | 0 | -0.100 | 0.080 | NA |
| SCFAs | Acetate degradation | Glutamate | DLPFC | 0.021 | 0.045 | 60 | 60 | 0.650 | 0.908 | 0.977 | 0 | -0.068 | 0.110 | NA |
| SCFAs | Acetate degradation | Glutamate | IOG | 0.042 | 0.047 | 60 | 60 | 0.379 | 0.774 | 0.948 | 0 | -0.050 | 0.135 | NA |
| SCFAs | Acetate synthesis I | GABA | ACC | -0.021 | 0.026 | 60 | 60 | 0.438 | 0.801 | 0.966 | 0 | -0.072 | 0.031 | NA |
| SCFAs | Acetate synthesis I | GABA | DLPFC | 0.040 | 0.022 | 60 | 60 | 0.085 | 0.616 | 0.601 | 3 | -0.004 | 0.083 | NA |
| SCFAs | Acetate synthesis I | GABA | IOG | 0.063 | 0.044 | 60 | 60 | 0.165 | 0.633 | 0.744 | 0 | -0.023 | 0.150 | NA |
| SCFAs | Acetate synthesis I | Glutamate | ACC | -0.012 | 0.027 | 60 | 60 | 0.647 | 0.904 | 0.976 | 0 | -0.065 | 0.040 | NA |
| SCFAs | Acetate synthesis I | Glutamate | DLPFC | -0.013 | 0.026 | 60 | 60 | 0.635 | 0.905 | 0.975 | 0 | -0.064 | 0.039 | NA |
| SCFAs | Acetate synthesis I | Glutamate | IOG | -0.023 | 0.027 | 60 | 60 | 0.421 | 0.788 | 0.955 | 0 | -0.076 | 0.031 | NA |
| SCFAs | Acetate synthesis II | GABA | ACC | 0.043 | 0.026 | 60 | 60 | 0.113 | 0.615 | 0.650 | 1 | -0.008 | 0.094 | NA |
| SCFAs | Acetate synthesis II | GABA | DLPFC | -0.029 | 0.022 | 60 | 60 | 0.204 | 0.643 | 0.796 | 1 | -0.072 | 0.014 | NA |
| SCFAs | Acetate synthesis II | GABA | IOG | 0.065 | 0.044 | 60 | 60 | 0.157 | 0.628 | 0.723 | 1 | -0.022 | 0.152 | NA |
| SCFAs | Acetate synthesis II | Glutamate | ACC | -0.047 | 0.027 | 60 | 60 | 0.090 | 0.617 | 0.615 | 2 | -0.099 | 0.006 | NA |
| SCFAs | Acetate synthesis II | Glutamate | DLPFC | 0.011 | 0.026 | 60 | 60 | 0.686 | 0.922 | 0.974 | 0 | -0.041 | 0.062 | NA |
| SCFAs | Acetate synthesis II | Glutamate | IOG | -0.017 | 0.027 | 60 | 60 | 0.530 | 0.853 | 0.973 | 0 | -0.071 | 0.036 | NA |
| SCFAs | Acetate synthesis III | GABA | ACC | 0.037 | 0.025 | 60 | 60 | 0.147 | 0.618 | 0.709 | 0 | -0.012 | 0.086 | NA |
| SCFAs | Acetate synthesis III | GABA | DLPFC | -0.029 | 0.021 | 60 | 60 | 0.174 | 0.628 | 0.752 | 1 | -0.070 | 0.012 | NA |
| SCFAs | Acetate synthesis III | GABA | IOG | 0.063 | 0.042 | 60 | 60 | 0.153 | 0.627 | 0.718 | 1 | -0.020 | 0.145 | NA |
| SCFAs | Acetate synthesis III | Glutamate | ACC | -0.043 | 0.025 | 60 | 60 | 0.099 | 0.616 | 0.631 | 1 | -0.092 | 0.007 | NA |
| SCFAs | Acetate synthesis III | Glutamate | DLPFC | 0.010 | 0.025 | 60 | 60 | 0.701 | 0.923 | 0.975 | 0 | -0.039 | 0.058 | NA |
| SCFAs | Acetate synthesis III | Glutamate | IOG | -0.015 | 0.026 | 60 | 60 | 0.574 | 0.881 | 0.975 | 0 | -0.066 | 0.036 | NA |
| SCFAs | Butyrate synthesis I | GABA | ACC | 0.076 | 0.073 | 60 | 60 | 0.311 | 0.724 | 0.918 | 0 | -0.067 | 0.219 | NA |
| SCFAs | Butyrate synthesis I | GABA | DLPFC | -0.084 | 0.061 | 60 | 60 | 0.187 | 0.633 | 0.766 | 1 | -0.203 | 0.036 | NA |
| SCFAs | Butyrate synthesis I | GABA | IOG | -0.026 | 0.123 | 60 | 60 | 0.802 | 0.941 | 0.969 | 0 | -0.267 | 0.215 | NA |
| SCFAs | Butyrate synthesis I | Glutamate | ACC | -0.102 | 0.074 | 60 | 60 | 0.184 | 0.639 | 0.771 | 1 | -0.247 | 0.044 | NA |
| SCFAs | Butyrate synthesis I | Glutamate | DLPFC | -0.077 | 0.073 | 60 | 60 | 0.304 | 0.727 | 0.917 | 0 | -0.221 | 0.066 | NA |
| SCFAs | Butyrate synthesis I | Glutamate | IOG | -0.084 | 0.076 | 60 | 60 | 0.285 | 0.720 | 0.903 | 0 | -0.233 | 0.066 | NA |
| SCFAs | Butyrate synthesis II | GABA | ACC | 0.270 | 0.560 | 60 | 57 | 0.632 | 0.905 | 0.976 | 0 | -0.827 | 1.368 | NA |
| SCFAs | Butyrate synthesis II | GABA | DLPFC | -0.107 | 0.469 | 60 | 57 | 0.815 | 0.952 | 0.977 | 0 | -1.026 | 0.812 | NA |
| SCFAs | Butyrate synthesis II | GABA | IOG | -0.158 | 0.943 | 60 | 57 | 0.864 | 0.965 | 0.977 | 0 | -2.005 | 1.690 | NA |
| **SCFAs** | **Butyrate synthesis II** | **Glutamate** | **ACC** | **-1.275** | **0.568** | **60** | **57** | **0.035** | **0.570** | **0.492** | **59** | **-2.389** | **-0.162** | **TRUE** |
| SCFAs | Butyrate synthesis II | Glutamate | DLPFC | -0.556 | 0.561 | 60 | 57 | 0.332 | 0.741 | 0.937 | 0 | -1.655 | 0.543 | NA |
| SCFAs | Butyrate synthesis II | Glutamate | IOG | -0.161 | 0.585 | 60 | 57 | 0.787 | 0.949 | 0.977 | 0 | -1.307 | 0.986 | NA |
| SCFAs | Propionate degradation I | GABA | ACC | -0.611 | 0.891 | 60 | 51 | 0.501 | 0.840 | 0.970 | 0 | -2.357 | 1.135 | NA |
| SCFAs | Propionate degradation I | GABA | DLPFC | 0.083 | 0.745 | 60 | 51 | 0.901 | 0.980 | 0.979 | 0 | -1.378 | 1.544 | NA |
| SCFAs | Propionate degradation I | GABA | IOG | -2.768 | 1.499 | 60 | 51 | 0.074 | 0.611 | 0.589 | 2 | -5.705 | 0.170 | NA |
| SCFAs | Propionate degradation I | Glutamate | ACC | 0.657 | 0.904 | 60 | 51 | 0.475 | 0.825 | 0.970 | 0 | -1.115 | 2.429 | NA |
| SCFAs | Propionate degradation I | Glutamate | DLPFC | -0.274 | 0.892 | 60 | 51 | 0.757 | 0.940 | 0.978 | 0 | -2.022 | 1.473 | NA |
| SCFAs | Propionate degradation I | Glutamate | IOG | 0.461 | 0.930 | 60 | 51 | 0.625 | 0.902 | 0.976 | 0 | -1.362 | 2.284 | NA |
| **SCFAs** | **Propionate synthesis III** | **GABA** | **ACC** | **0.223** | **0.101** | **60** | **60** | **0.035** | **0.573** | **0.498** | **55** | **0.025** | **0.420** | **TRUE** |
| SCFAs | Propionate synthesis III | GABA | DLPFC | -0.127 | 0.084 | 60 | 60 | 0.154 | 0.623 | 0.696 | 1 | -0.292 | 0.038 | NA |
| SCFAs | Propionate synthesis III | GABA | IOG | -0.373 | 0.170 | 60 | 60 | 0.041 | 0.581 | 0.508 | 57 | -0.706 | -0.041 | TRUE |
| SCFAs | Propionate synthesis III | Glutamate | ACC | -0.165 | 0.102 | 60 | 60 | 0.120 | 0.622 | 0.669 | 0 | -0.365 | 0.036 | NA |
| SCFAs | Propionate synthesis III | Glutamate | DLPFC | 0.065 | 0.101 | 60 | 60 | 0.507 | 0.836 | 0.971 | 0 | -0.133 | 0.263 | NA |
| SCFAs | Propionate synthesis III | Glutamate | IOG | 0.094 | 0.105 | 60 | 60 | 0.381 | 0.771 | 0.949 | 0 | -0.112 | 0.300 | NA |

### Associations between Neuroactive Pathways in the microbiota and Brain Neurotransmitter Levels

#### Microbial Taxonomic Contributors to Neuroactive Pathways

Functional profiling of gut microbiota identified key microbial taxa associated with the neuroactive pathways evaluated. GABA synthesis pathways (GABA synthesis I and II) were predominantly contributed by Bifidobacterium species (e.g., *B. adolescentis*, *B. dentium*) and Lactobacillus species (e.g., *L. brevis*, *L. rhamnosus*), consistent with prior evidence of their psychobiotic potential [4, 69, 70]. The p-cresol synthesis pathway was mainly attributed to Clostridium species, particularly members of the *Clostridium cluster XI*, known for aromatic compound metabolism [16, 71]. Short-chain fatty acid (SCFA) synthesis pathways were largely associated with *Faecalibacterium prausnitzii*, *Roseburia*, and *Eubacterium* species, reflecting established SCFA-producing lineages. These taxa were mapped onto the neuroactive pathway models using gene-function annotations from the KEGG ORTHOLOGY database. These microbial contributors form the basis for the neuroactive pathways analyzed below, where we report their associations with brain GABA and glutamate concentrations across the dlPFC, ACC, and IOG.

#### Amino Acid Metabolism

Analysis revealed several significant region-specific associations (Figure 2). In the dorsolateral prefrontal cortex (dlPFC), glutamate levels negatively predicted activity in the gut microbial GABA synthesis I pathway (β = −3.482, p = 0.008, lfdr = 0.380, 95% CI [−5.896, −1.068]). Additionally, in the inferior occipital gyrus (IOG), GABA levels negatively correlated with the GABA synthesis II pathway (β = −4.749, p = 0.034, lfdr = 0.497, 95% CI [−8.963, - 0.535]).

**Figure 2.**
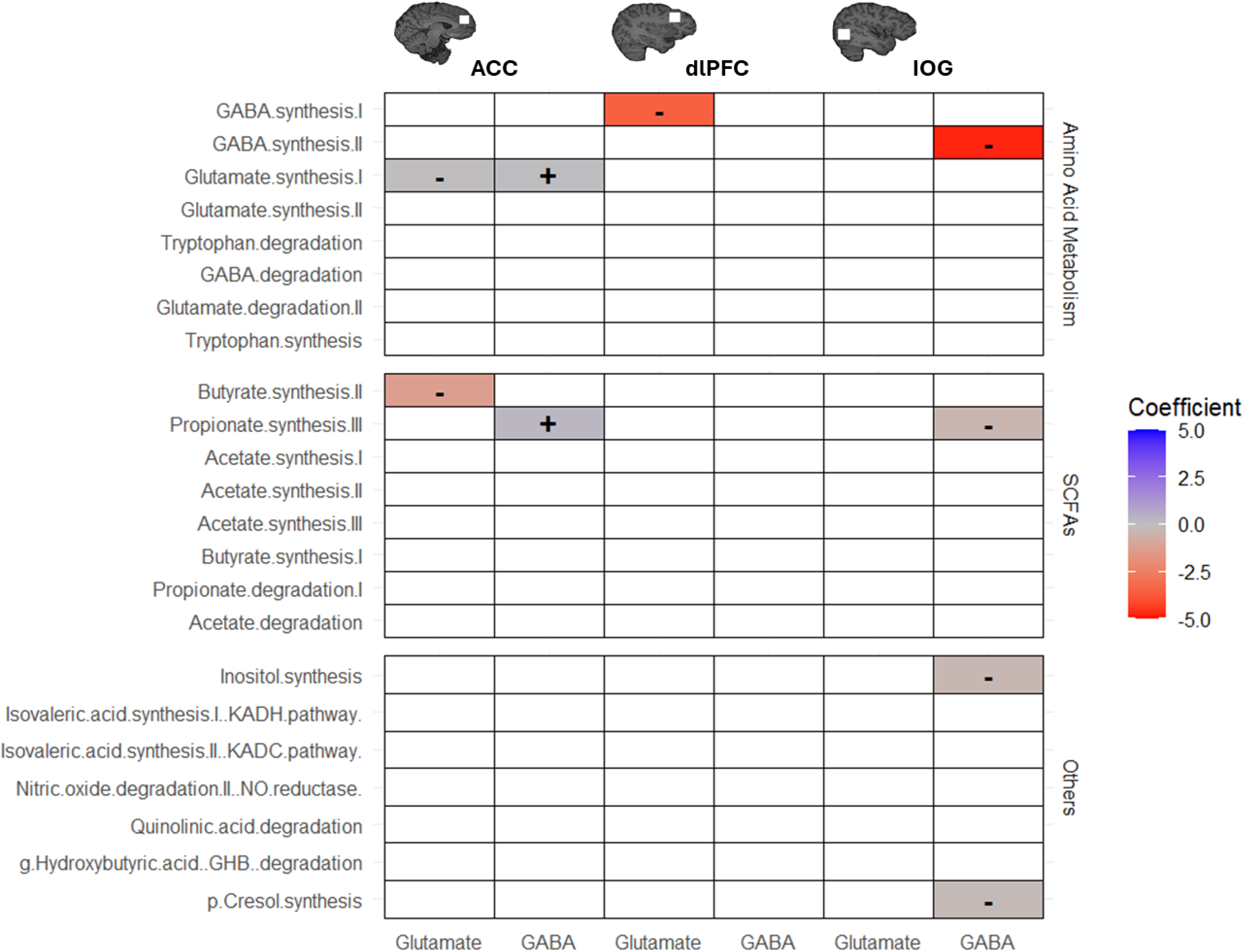
Heatmap of model coefficients of neuroactive pathways in the gut and brain glutamate and GABA levels in different regions. Magnitude and direction of coefficient signified by colour. Only effects with a 95% confidence interval that does not cross zero are considered true effects and plotted.

For the glutamate synthesis I pathway, two significant associations were observed in the anterior cingulate cortex (ACC): GABA levels positively predicted pathway activity (β = 0.042, p = 0.021, lfdr = 0.452, 95% CI [0.008, 0.075]), while glutamate exhibited a negative association (β = −0.036, p = 0.050, lfdr = 0.533, 95% CI [−0.070, −0.001]).

#### Short-Chain Fatty Acid (SCFA) Synthesis

Butyrate synthesis II pathway activity negatively correlated with glutamate levels in the ACC (β = −1.275, p = 0.035, lfdr = 0.492, 95% CI [−2.389, −0.162]). Additionally, propionate synthesis III exhibited a significant positive association with GABA levels in the ACC (β = 0.223, p = 0.035, lfdr = 0.498, 95% CI [0.025, 0.420]) and a negative association in the IOG (β = −0.373, p = 0.041, lfdr = 0.508, 95% CI [−0.706, −0.041]).

#### Other Neuroactive Pathways

In the IOG, GABA levels negatively correlated with the inositol synthesis pathway (β = - 0.317, p = 0.006, lfdr = 0.366, 95% CI [−0.531, −0.102]). Similarly, GABA levels in the IOG showed a negative association with the p-cresol synthesis pathway (β = −0.174, p = 0.015, lfdr = 0.423, 95% CI [−0.305, −0.043]). Full model results are presented in **Table 1**.

#### Associations between Neuroactive Pathways and E/I Ratio

Analysis of neuroactive pathways and their association with the excitatory/inhibitory (E/I) ratio identified nine significant associations, two in the anterior cingulate cortex (ACC) and seven in the inferior occipital gyrus (IOG) **Figure 3**.

**Figure 3.**
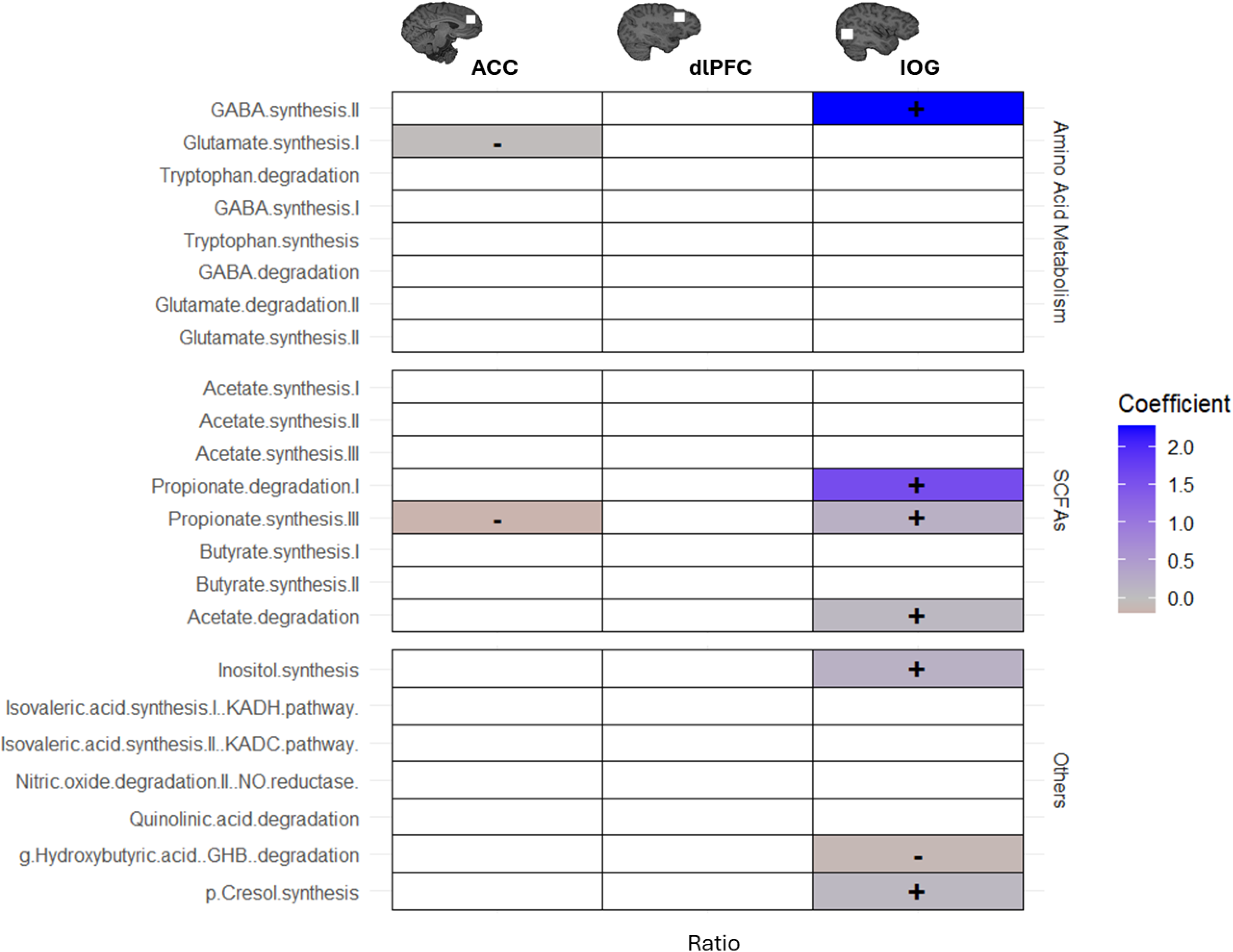
Heatmap of model coefficients of neuroactive pathways in the gut and brain E/I ratios in different regions. Only effects with a 95% confidence interval that does not cross zero are considered true effects and plotted.

#### Amino Acid Metabolism

In the ACC, the glutamate synthesis I pathway negatively correlated with the E/I ratio (β = - 0.032, p = 0.029, lfdr = 0.308, 95% CI [−0.059, −0.004]). In contrast, the GABA synthesis II pathway in the IOG positively correlated with the E/I ratio (β = 2.266, p = 0.050, lfdr = 0.392, 95% CI [0.077, 4.456]).

#### Short-Chain Fatty Acids (SCFAs)

The acetate degradation pathway exhibited a significant positive relationship with the E/I ratio in the IOG (β = 0.084, p = 0.034, lfdr = 0.330, 95% CI [0.010, 0.158]). Propionate synthesis III pathway showed region-specific relationships with the E/I ratio: negatively in the ACC (β = −0.197, p = 0.020, lfdr = 0.254, 95% CI [−0.352, −0.041]) and positively in the IOG (β = 0.193, p = 0.032, lfdr = 0.322, 95% CI [0.024, 0.362]). Additionally, propionate degradation I pathway positively correlated with the E/I ratio in the IOG (β = 1.577, p = 0.043, lfdr = 0.358, 95% CI [0.111, 3.042]).

#### Other Neuroactive Pathways

The p-cresol synthesis pathway exhibited a positive association with the E/I ratio in the IOG (β = 0.083, p = 0.022, lfdr = 0.267, 95% CI [0.015, 0.151]). Conversely, g-Hydroxybutyric acid (GHB) degradation showed a negative association within the same region (β = −0.113, p = 0.053, lfdr = 0.406, 95% CI [−0.223, −0.002]). Finally, inositol synthesis positively correlated with the E/I ratio in the IOG (β = 0.187, p = 0.002, lfdr = 0.101, 95% CI [0.079, 0.294]). Full model results are reported in **Table 2**.

**Table 2.**
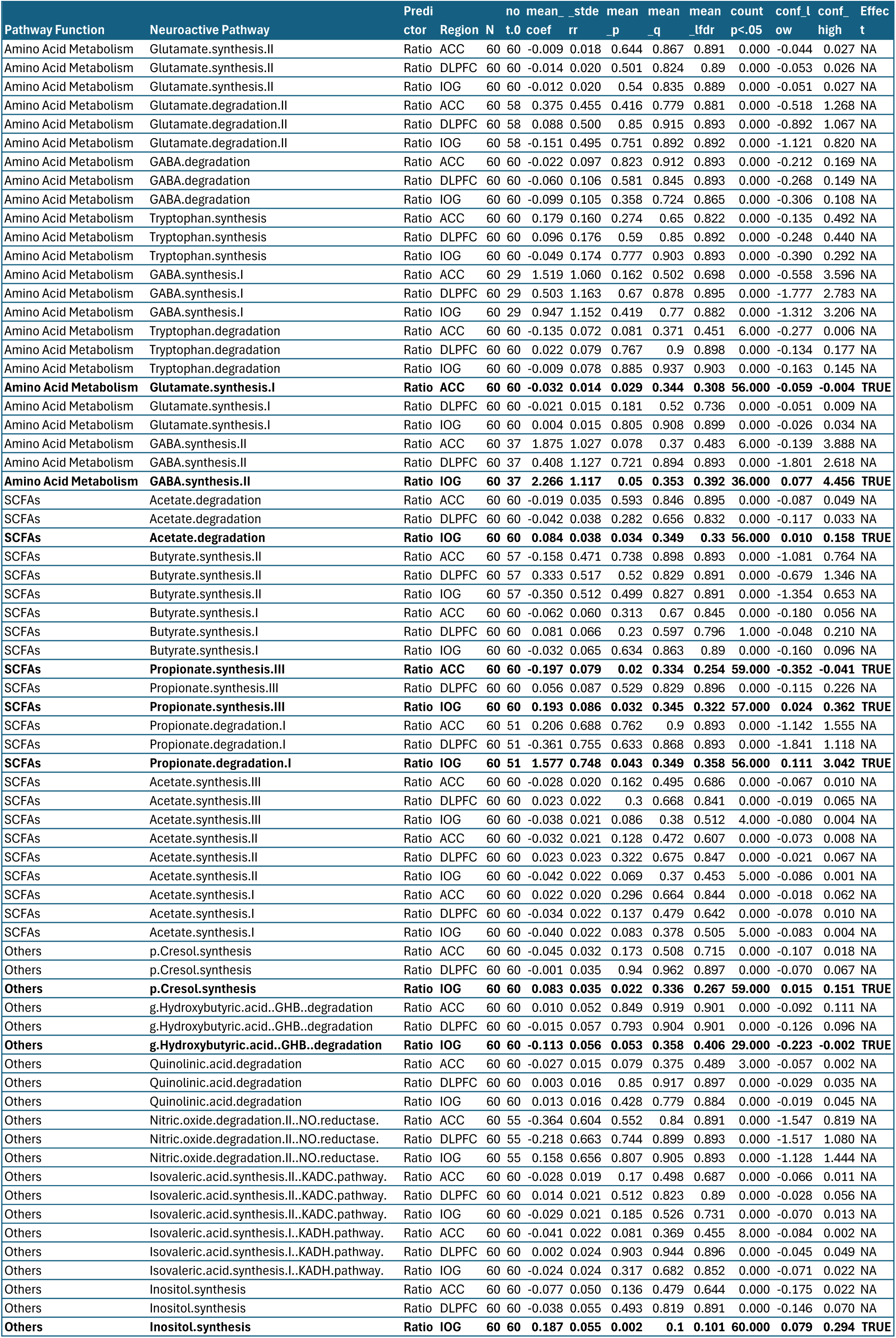
Strength and direction of relationship between the E/I ratio by region and neuroactive pathways of gut microbiota.

**Table 3.** Strength and direction of relationship between psychological varibles and neuroactive pathways of gut microbiota.

| Pathway function | Neuroactive Pathway | Predictor | coef | stderr | N | N.not.0 | pval | qval | lfdr |
| --- | --- | --- | --- | --- | --- | --- | --- | --- | --- |
| **Amino Acid Metabolism** | **GABA degradation** | **Trait** | **-0.386** | **0.183** | **61.000** | **61.000** | **0.040** | **0.668** | **0.503** |
| Amino Acid Metabolism | GABA degradation | PSQI | 0.131 | 0.137 | 61.000 | 61.000 | 0.343 | 0.696 | 0.626 |
| Amino Acid Metabolism | GABA degradation | BDI | 0.157 | 0.189 | 61.000 | 61.000 | 0.411 | 0.720 | 0.674 |
| **Amino Acid Metabolism** | **GABA synthesis I** | **Trait** | **0.610** | **0.497** | **61.000** | **29.000** | **0.225** | **0.668** | **0.544** |
| **Amino Acid Metabolism** | **GABA synthesis I** | **BDI** | **-0.686** | **0.514** | **61.000** | **29.000** | **0.187** | **0.668** | **0.519** |
| Amino Acid Metabolism | GABA synthesis I | PSQI | -0.089 | 0.372 | 61.000 | 29.000 | 0.813 | 0.878 | 1.000 |
| **Amino Acid Metabolism** | **GABA synthesis II** | **BDI** | **-1.101** | **0.588** | **61.000** | **38.000** | **0.066** | **0.668** | **0.503** |
| Amino Acid Metabolism | GABA synthesis II | Trait | 0.555 | 0.569 | 61.000 | 38.000 | 0.333 | 0.696 | 0.619 |
| Amino Acid Metabolism | GABA synthesis II | PSQI | 0.141 | 0.426 | 61.000 | 38.000 | 0.742 | 0.840 | 1.000 |
| **Amino Acid Metabolism** | **Glutamate degradation II** | **Trait** | **-0.682** | **0.508** | **61.000** | **59.000** | **0.185** | **0.668** | **0.518** |
| **Amino Acid Metabolism** | **Glutamate degradation II** | **BDI** | **0.719** | **0.525** | **61.000** | **59.000** | **0.177** | **0.668** | **0.512** |
| **Amino Acid Metabolism** | **Glutamate degradation II** | **PSQI** | **-0.640** | **0.380** | **61.000** | **59.000** | **0.098** | **0.668** | **0.503** |
| Amino Acid Metabolism | Glutamate synthesis I | PSQI | 0.020 | 0.021 | 61.000 | 61.000 | 0.331 | 0.696 | 0.617 |
| Amino Acid Metabolism | Glutamate synthesis I | Trait | 0.017 | 0.028 | 61.000 | 61.000 | 0.545 | 0.720 | 0.791 |
| Amino Acid Metabolism | Glutamate synthesis I | BDI | -0.021 | 0.029 | 61.000 | 61.000 | 0.475 | 0.720 | 0.724 |
| **Amino Acid Metabolism** | **Glutamate synthesis II** | **PSQI** | **-0.046** | **0.027** | **61.000** | **61.000** | **0.098** | **0.668** | **0.503** |
| Amino Acid Metabolism | Glutamate synthesis II | Trait | 0.038 | 0.036 | 61.000 | 61.000 | 0.297 | 0.696 | 0.594 |
| Amino Acid Metabolism | Glutamate synthesis II | BDI | 0.012 | 0.037 | 61.000 | 61.000 | 0.743 | 0.840 | 1.000 |
| **Amino Acid Metabolism** | **Tryptophan degradation** | **BDI** | **-0.205** | **0.148** | **61.000** | **61.000** | **0.173** | **0.668** | **0.510** |
| **Amino Acid Metabolism** | **Tryptophan degradation** | **PSQI** | **0.130** | **0.108** | **61.000** | **61.000** | **0.231** | **0.668** | **0.548** |
| Amino Acid Metabolism | Tryptophan degradation | Trait | 0.083 | 0.144 | 61.000 | 61.000 | 0.565 | 0.720 | 0.813 |
| **Amino Acid Metabolism** | **Tryptophan synthesis** | **PSQI** | **0.329** | **0.220** | **61.000** | **61.000** | **0.140** | **0.668** | **0.503** |
| Amino Acid Metabolism | Tryptophan synthesis | BDI | -0.300 | 0.303 | 61.000 | 61.000 | 0.327 | 0.696 | 0.614 |
| Amino Acid Metabolism | Tryptophan synthesis | Trait | -0.129 | 0.293 | 61.000 | 61.000 | 0.663 | 0.803 | 0.955 |
| Aromatic Compound Metabolism | Quinolinic acid degradation | BDI | 0.031 | 0.031 | 61.000 | 61.000 | 0.320 | 0.696 | 0.610 |
| Aromatic Compound Metabolism | Quinolinic acid degradation | PSQI | 0.014 | 0.022 | 61.000 | 61.000 | 0.527 | 0.720 | 0.772 |
| Aromatic Compound Metabolism | Quinolinic acid degradation | Trait | -0.002 | 0.030 | 61.000 | 61.000 | 0.955 | 0.976 | 1.000 |
| **Aromatic Compound Metabolism** | **p Cresol synthesis** | **Trait** | **0.087** | **0.060** | **61.000** | **61.000** | **0.153** | **0.668** | **0.503** |
| Aromatic Compound Metabolism | p Cresol synthesis | BDI | -0.073 | 0.062 | 61.000 | 61.000 | 0.248 | 0.668 | 0.560 |
| Aromatic Compound Metabolism | p Cresol synthesis | PSQI | 0.011 | 0.045 | 61.000 | 61.000 | 0.814 | 0.878 | 1.000 |
| **Inositol Metabolism** | **Inositol synthesis** | **Trait** | **0.160** | **0.099** | **61.000** | **61.000** | **0.113** | **0.668** | **0.503** |
| **Inositol Metabolism** | **Inositol synthesis** | **BDI** | **-0.187** | **0.103** | **61.000** | **61.000** | **0.074** | **0.668** | **0.503** |
| Inositol Metabolism | Inositol synthesis | PSQI | 0.089 | 0.074 | 61.000 | 61.000 | 0.235 | 0.668 | 0.551 |
| Nitrogen Cycle | Nitric oxide degradation II NO reductas | Trait | -0.549 | 0.474 | 61.000 | 56.000 | 0.252 | 0.668 | 0.562 |
| Nitrogen Cycle | Nitric oxide degradation II NO reductas | PSQI | -0.327 | 0.355 | 61.000 | 56.000 | 0.360 | 0.710 | 0.638 |
| Nitrogen Cycle | Nitric oxide degradation II NO reductas | BDI | 0.319 | 0.490 | 61.000 | 56.000 | 0.518 | 0.720 | 0.763 |
| Organic Acids and Other Pathways | Isovaleric acid synthesis I KADH pathw | PSQI | 0.028 | 0.035 | 61.000 | 61.000 | 0.419 | 0.720 | 0.680 |
| Organic Acids and Other Pathways | Isovaleric acid synthesis I KADH pathw | Trait | -0.024 | 0.047 | 61.000 | 61.000 | 0.603 | 0.743 | 0.860 |
| Organic Acids and Other Pathways | Isovaleric acid synthesis I KADH pathw | BDI | -0.002 | 0.048 | 61.000 | 61.000 | 0.962 | 0.976 | 1.000 |
| Organic Acids and Other Pathways | Isovaleric acid synthesis II KADC pathw | Trait | -0.036 | 0.041 | 61.000 | 61.000 | 0.391 | 0.720 | 0.660 |
| Organic Acids and Other Pathways | Isovaleric acid synthesis II KADC pathw | BDI | 0.032 | 0.043 | 61.000 | 61.000 | 0.459 | 0.720 | 0.711 |
| Organic Acids and Other Pathways | Isovaleric acid synthesis II KADC pathw | PSQI | 0.000 | 0.031 | 61.000 | 61.000 | 0.993 | 0.993 | 1.000 |
| **Organic Acids and Other Pathways** | **g Hydroxybutyric acid GHB degradati** | **Trait** | **-0.209** | **0.115** | **61.000** | **61.000** | **0.075** | **0.668** | **0.503** |
| **Organic Acids and Other Pathways** | **g Hydroxybutyric acid GHB degradati** | **PSQI** | **0.107** | **0.086** | **61.000** | **61.000** | **0.220** | **0.668** | **0.541** |
| Organic Acids and Other Pathways | g Hydroxybutyric acid GHB degradati | BDI | 0.073 | 0.119 | 61.000 | 61.000 | 0.538 | 0.720 | 0.783 |
| SCFAs | Acetate degradation | Trait | 0.055 | 0.068 | 61.000 | 61.000 | 0.425 | 0.720 | 0.684 |
| SCFAs | Acetate degradation | BDI | -0.048 | 0.070 | 61.000 | 61.000 | 0.500 | 0.720 | 0.746 |
| SCFAs | Acetate degradation | PSQI | 0.029 | 0.051 | 61.000 | 61.000 | 0.566 | 0.720 | 0.813 |
| **SCFAs** | **Acetate synthesis I** | **BDI** | **0.064** | **0.043** | **61.000** | **61.000** | **0.140** | **0.668** | **0.503** |
| SCFAs | Acetate synthesis I | Trait | -0.047 | 0.042 | 61.000 | 61.000 | 0.262 | 0.668 | 0.569 |
| SCFAs | Acetate synthesis I | PSQI | -0.028 | 0.031 | 61.000 | 61.000 | 0.373 | 0.716 | 0.647 |
| SCFAs | Acetate synthesis II | PSQI | 0.036 | 0.030 | 61.000 | 61.000 | 0.240 | 0.668 | 0.554 |
| SCFAs | Acetate synthesis II | Trait | -0.028 | 0.040 | 61.000 | 61.000 | 0.483 | 0.720 | 0.731 |
| SCFAs | Acetate synthesis II | BDI | 0.024 | 0.041 | 61.000 | 61.000 | 0.568 | 0.720 | 0.816 |
| **SCFAs** | **Acetate synthesis III** | **PSQI** | **0.037** | **0.028** | **61.000** | **61.000** | **0.204** | **0.668** | **0.530** |
| SCFAs | Acetate synthesis III | Trait | -0.026 | 0.038 | 61.000 | 61.000 | 0.490 | 0.720 | 0.737 |
| SCFAs | Acetate synthesis III | BDI | 0.022 | 0.039 | 61.000 | 61.000 | 0.574 | 0.720 | 0.823 |
| SCFAs | Butyrate synthesis I | PSQI | 0.093 | 0.088 | 61.000 | 61.000 | 0.294 | 0.696 | 0.592 |
| SCFAs | Butyrate synthesis I | Trait | -0.074 | 0.117 | 61.000 | 61.000 | 0.532 | 0.720 | 0.777 |
| SCFAs | Butyrate synthesis I | BDI | 0.007 | 0.121 | 61.000 | 61.000 | 0.956 | 0.976 | 1.000 |
| **SCFAs** | **Butyrate synthesis II** | **BDI** | **-0.700** | **0.443** | **61.000** | **58.000** | **0.120** | **0.668** | **0.503** |
| **SCFAs** | **Butyrate synthesis II** | **PSQI** | **0.617** | **0.321** | **61.000** | **58.000** | **0.060** | **0.668** | **0.503** |
| SCFAs | Butyrate synthesis II | Trait | 0.166 | 0.429 | 61.000 | 58.000 | 0.700 | 0.832 | 1.000 |
| **SCFAs** | **Propionate degradation I** | **BDI** | **-0.572** | **0.434** | **61.000** | **52.000** | **0.193** | **0.668** | **0.522** |
| SCFAs | Propionate degradation I | Trait | 0.121 | 0.420 | 61.000 | 52.000 | 0.774 | 0.862 | 1.000 |
| SCFAs | Propionate degradation I | PSQI | -0.034 | 0.314 | 61.000 | 52.000 | 0.915 | 0.971 | 1.000 |
| **SCFAs** | **Propionate synthesis III** | **PSQI** | **0.232** | **0.126** | **61.000** | **61.000** | **0.072** | **0.668** | **0.503** |
| SCFAs | Propionate synthesis III | BDI | -0.101 | 0.174 | 61.000 | 61.000 | 0.564 | 0.720 | 0.812 |
| SCFAs | Propionate synthesis III | Trait | 0.058 | 0.169 | 61.000 | 61.000 | 0.733 | 0.840 | 1.000 |

#### Associations Between Microbial Pathways and Psychological Outcomes

To explore the potential psychological relevance of the observed gut-brain associations, we examined relationships between microbial neuroactive pathways and key self-reported psychological outcomes, including trait anxiety, sleep quality, and depressive symptoms.

These associations are summarized in Figure 4, which highlights specific microbial pathways showing significant links to psychological traits. Several microbial pathways previously associated with neurotransmitter balance also showed associations with psychological outcomes (**Figure 4**). The GABA degradation pathway exhibited a negative association with trait anxiety (β = –0.386, p = 0.040, lfdr = 0.503, 95% CI [–0.747, –0.025]). The GABA synthesis II pathway was negatively associated with depressive symptoms (β = –1.101, p = 0.066, lfdr = 0.503, 95% CI [–2.254, 0.051]).

**Figure 4.**
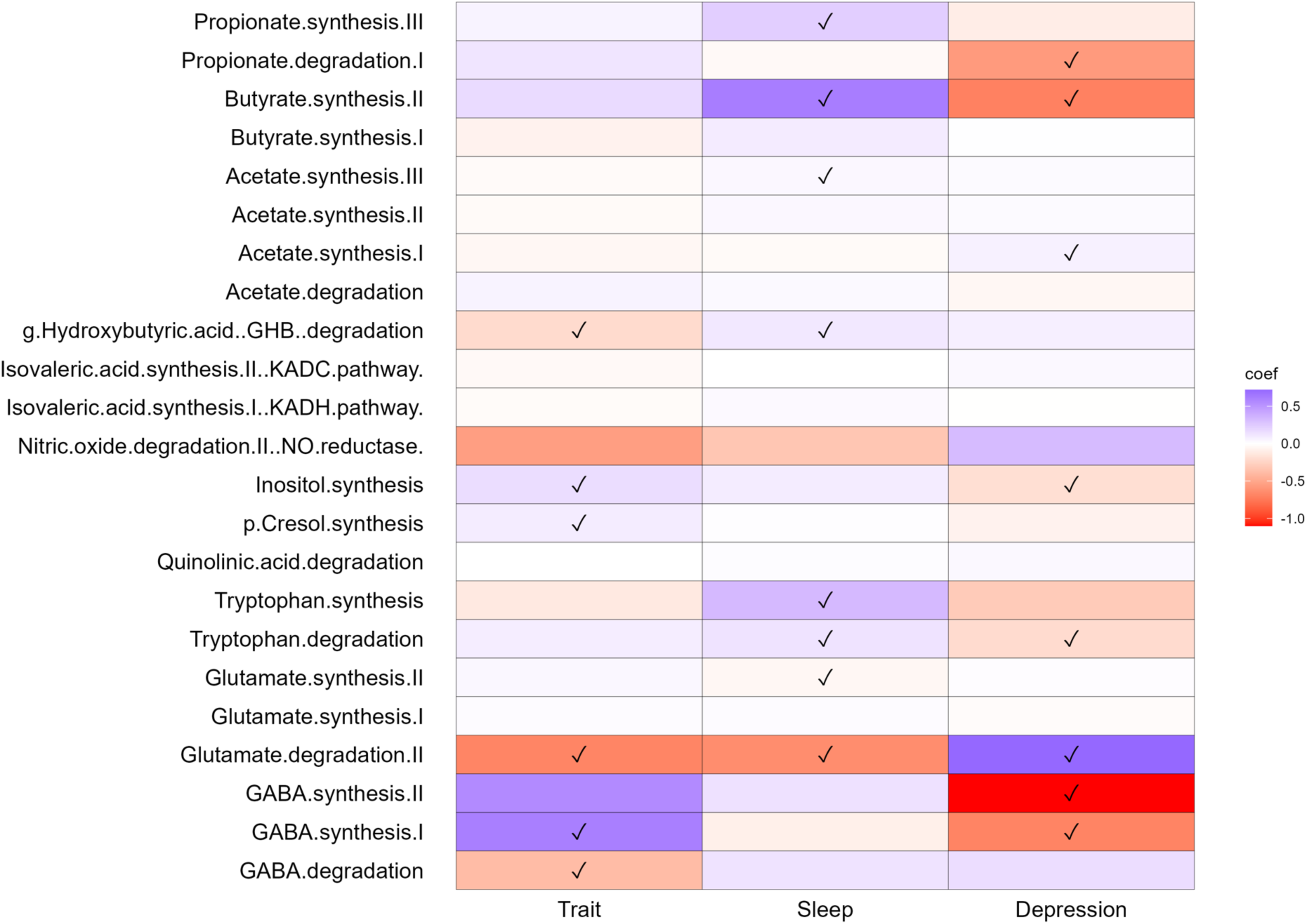
Associations between microbial neuroactive pathways and psychological outcomes. Heatmap displaying the direction and strength of associations (coefficients) between microbial pathway activity and trait anxiety, sleep quality, and depressive symptoms. Pathways highlighted include those involved in neurotransmitter metabolism (GABA, glutamate), aromatic compound degradation (p-cresol), and SCFA synthesis. Darker shading indicates stronger associations (black = positive, purple = negative; local false discover rate <50 lfdr).

For glutamate-related pathways, glutamate degradation II was positively associated with depressive symptoms (β = 0.719, p = 0.177, lfdr = 0.512, 95% CI [–0.312, 1.750]) and negatively associated with trait anxiety (β = –0.682, p = 0.185, lfdr = 0.518, 95% CI [–1.687, 0.322]). The p-cresol synthesis pathway showed a positive association with trait anxiety (β = 0.087, p = 0.153, lfdr = 0.503, 95% CI [–0.032, 0.207]). Full model results are reported in F**igure 4**.

## Discussion

Consistent with our hypotheses, our results suggest that gut-derived neuroactive pathways influence brain neurotransmitter dynamics, E/I balance, and psychological functioning. Region-specific associations between microbial metabolism, GABA and glutamate levels, and E/I ratios were observed across the anterior cingulate cortex (ACC), dorsolateral prefrontal cortex (dlPFC), and inferior occipital gyrus (IOG), with several pathways also showing links to mood, anxiety, and sleep quality. Below, we discuss these converging findings in relation to brain function.

In the ACC, microbial glutamate synthesis I was associated with increased GABA concentrations and reduced E/I balance, suggesting that gut-derived glutamate may support inhibitory tone through GABAergic pathways. Glutamate is a critical energy source within the gut, where it is converted from α-ketoglutarate and ammonia by glutamate dehydrogenase (GDH) [72]. Glutamate, in turn, serves as a precursor to GABA, supporting the observed relationship between microbial glutamate synthesis and brain GABA levels in this region [73]. However, this association was only detected in the ACC and not in the inferior occipital gyrus (IOG) or the dorsolateral prefrontal cortex (dlPFC). The ACC supports both attention and executive function, as well as emotion regulation and the processing of fear and stress. Critically, research has shown altered ACC responsiveness in individuals with anxiety [74–76]. This specificity suggests additional regional factors, such as receptor distribution, local metabolic demands, or distinct vulnerability to microbial metabolites. The observed convergence between neurotransmitter concentrations, E/I ratio, and microbial pathway activity within the ACC supports the concept of microbial modulation of excitatory tone and cortical inhibition in this region. Our results also support the concept of targeted therapeutic strategies based on the understanding of highly localized gut-brain interactions [18].

In the dlPFC, a strong negative relationship emerged between the GABA synthesis I pathway and brain glutamate levels. Microbial species involved in GABA synthesis, such as strains of bifidobacteria (e.g., *B. adolescentis*, *B. dentium*) and *Lactobacillus* species (e.g., L*. brevis, L. rhamnosus*)[69, 70], were found to influence this pathway. This relationship suggests that microbial GABA synthesis may regulate cortical excitability via glutamatergic pathways. This is particularly relevant for the dlPFC, a region critical for emotion regulation, cognitive control, and executive functioning [41, 42]. Although E/I ratios did not show significant associations in the dlPFC, the neurotransmitter findings implicate this region as another site where microbial GABA pathways may influence prefrontal function, complementing the effects observed in the ACC.

In the IOG, brain GABA levels showed a weak negative association with the synthesis of inositol and p-cresol in the gut. Inositol, a secondary messenger critical for intracellular signalling, does not cross the blood-brain barrier (BBB), so its relationship with brain GABA is likely indirect. In contrast, p-cresol, a neurotoxic compound, has been implicated in various neurological disorders, including autism spectrum disorder, and may contribute to increased brain excitability [16]. Previous studies have suggested that high p-cresol concentrations can disrupt brain dopamine metabolism, further exacerbating neurodevelopmental conditions.

However, emerging evidence indicates that low doses of p-cresol may exert distinct effects, potentially influencing neuronal metabolism and synapse remodelling. Building on previous work that links p-cresol to oxidative stress and downstream brain-derived neurotrophic factor (BDNF) signalling, a critical player in neuroplasticity [71], we propose that this microbial metabolite might influence neuroplasticity via similar pathways. Although we did not measure oxidative stress or BDNF directly in the present study, this raises the possibility that specific gut microbiome strains, particularly Clostridium species, and their metabolites might modulate neuroplasticity and brain function in both adaptive and maladaptive ways. The associations identified between GABA and both inositol and p-cresol synthesis pathways in the IOG suggest indirect microbial pathways might modulate cortical neurotransmission.

These findings were further supported by E/I ratio analyses, which highlighted the IOG as a region of prominent microbial influence. Pathways including GABA synthesis II, acetate degradation, propionate synthesis and degradation, and p-cresol synthesis exhibited significant associations with E/I balance in the IOG (Figure 3). This suggests that microbial metabolic activity may modulate cortical excitability in visual regions, raising questions about the broader functional impact of these pathways beyond prefrontal areas. Taken together, these findings highlight distinct region-specific signatures through which gut-derived metabolites may shape neurotransmitter balance, cortical excitability, and psychological function.

Several mechanistic routes may underlie these effects, including vagal nerve signalling, immune modulation, and microbial influence on barrier integrity and systemic metabolite availability. Short-chain fatty acids, GABA, and p-cresol emerged as key microbial pathways linked to brain neurochemistry and psychological outcomes in this study, underscoring their relevance as candidate targets for microbiome-based interventions. While GABA is traditionally regarded as an inhibitory neurotransmitter, its excitatory effects under certain neurochemical conditions—particularly during neurodevelopment—highlight the need to consider broader physiological contexts when interpreting gut-brain interactions [77].

Although ¹H-MRS provides estimates of neurotransmitter concentrations in vivo, it remains a proxy for true neurotransmission. Future research integrating ¹H-MRS with complementary approaches such as electrophysiology, functional imaging, and metabolomics will be critical to further elucidate the pathways through which gut microbial metabolites influence brain activity. Longitudinal and interventional studies will also be essential to assess causality and determine whether modulation of these microbial pathways can enhance cognitive resilience and mental wellbeing. Central to this work is the modulation of excitatory/inhibitory (E/I) balance—the equilibrium between excitatory and inhibitory neurotransmission—which represents a critical metric for neuroplasticity, mental health, and cognition [9, 11–13].

Emerging research on psychobiotics—including prebiotics and probiotics that influence gut microbiota composition—offers promising therapeutic potential for targeting the gut-brain axis [78].

### Study limitations

A limitation of the current study is the interpretative complexity introduced by regional variability in the brain. Future studies should integrate microbiome data, neurotransmitter receptor profiling, and additional neuroimaging modalities to further elucidate these relationships. In line with recent recommendations by Dalile et al. [79], broadening the scope of future research to include diverse populations and challenging conditions—whether in daily life or experimental settings—will be crucial for expanding our understanding of microbiome–brain interactions and for identifying pathways that may promote cognitive resilience and mental wellbeing.

In addition, several physiological and methodological factors may confound the associations observed here. For example, we did not account for individual circadian rhythms that may influence neuroimaging and stool sample measurements that are known to exhibit diurnal fluctuations. Menstrual cycle phase is also not controlled and may influence both neurotransmitter dynamics and gut microbiome composition, particularly in female participants. Similarly, this single sex sample may not be representative of male participants given neurochemical variations and interacting hormonal influences between male and female sexes.

This study contributes foundational knowledge to this growing field, demonstrating integrative links between microbial metabolism, brain neurochemistry, and psychological health. In particular, the observed associations between microbial pathways, neurotransmitter dynamics, and psychological outcomes represent a novel finding that bridges molecular, neural, and experiential levels of analysis, providing a rationale for microbiome-targeted strategies to support emotional health, cognitive function, and stress resilience. Several mechanisms could underlie these effects, including vagal signalling, direct metabolite action on peripheral neurons, immune modulation, or microbial influence on barrier integrity and systemic metabolite availability. Short-chain fatty acids, GABA, and p-cresol emerged as key microbial pathways linked to brain neurochemistry and psychological outcomes in this study, underscoring their relevance as candidate targets for future microbiome-based interventions.

## Acknowledgements

We thank Melissa Basso, Eloise Crowson, Nathalie Dusart, Hannah Eiseman, Chloe Hambleton, Daniella Louise Jones, Paul Knytl, Mariia Malova, and Greta Tomasi for supporting the production of research materials and data in this study.

## Data availability

The data that support the findings of this study are openly available in osf at **TBC.**

## CRediT Authorship

**Nicola Johnstone**: conceptualisation, data curation (lead), formal analysis (lead), funding acquisition, investigation (equal), methodology (equal), project administration (equal), visualisation (lead), writing – original draft preparation (lead), writing – review and editing (equal). **Kathrin Cohen Kadosh**: conceptualisation, funding acquisition (lead), investigation (equal), methodology (equal), project administration (equal), writing – review and editing (equal).

